# A global map for introgressed structural variation and selection in humans

**DOI:** 10.1101/2025.06.24.661368

**Authors:** PingHsun Hsieh, Natthapon Soisangwan, David S. Gordon, Athef Javidh, William T. Harvey, David Porubsky, Kendra Hoekzema, Carl Baker, Katherine M. Munson, Christopher Kinipi, Matthew Leavesley, Nicolas Brucato, Murray P. Cox, François-X Ricaut, Irene Gallego Romero, Evan E. Eichler

## Abstract

Genetic introgression from Neanderthals and Denisovan has shaped modern human genomes; however, introgressed structural variants (SVs ≥50 base pairs) remain challenging to discover. We integrated high-quality phased assemblies from four new Papua New Guinea (PNG) genomes with 94 published assemblies of diverse ancestry to infer an archaic introgressed SV map. Introgressed SVs are overall enriched in genes (44%, n=1,592), including critical genomic disorder regions, and most abundant in PNG. We identify 11 centromeres likely derived from archaic hominins, adding unexplored diversity to centromere genomics. Pangenome genotyping across 1,363 samples reveals 16 candidate adaptive SVs, many associated with immune-related genes and their expression, in the PNG. We hypothesize that archaic SV introgression contributed to reproductive success, underscoring introgression as a significant force in human adaptive evolution.

## INTRODUCTION

Evidence from over a decade of research unequivocally supports interbreeding occurred between archaic hominins, such as Neanderthals(*1–3*) and Denisovan(*4*), and the ancestors of modern humans, likely through multiple points of contact over the past 100,000 years of human evolution(*5, 6*). Genomic studies have largely focused on using single-nucleotide variant (SNV) from diverse populations to establish patterns of archaic introgression in our genome. Modern humans in Eurasia today derive 2–5% (or 120– 300 million base pairs [Mbp] per diploid genome) of their ancestry from archaic hominins, with the highest levels observed in Papua New Guinea (PNG)(*5, 7*). Despite the evidence of selection against deleterious archaic alleles in the human genome, some archaic sequences likely contributed to human phenotypic variation(*8, 9*). Consistent with these functional implications, many introgressed loci in our genome show signatures of positive selection. Some of those loci encompass candidate genes that are known to be functionally associated with differential gene expression, altitude, immunity, metabolism, and disease, highlighting contributions of introgressed alleles to the evolution of our species(*10–16*). Notwithstanding these discoveries, our understanding of the contribution of archaic introgression to human genomic variation and evolution remains far from complete due to the incomplete characterization of all classes of genetic variation. Structural variants (SVs), especially in complex loci, including segmental duplications (SDs) and centromeres(*17–19*), have been challenging to assess because of the highly fragmented nature of ancient DNA (∼50 bp) and the near impossibility of systematically discovering SVs from such ancient DNA, especially in gene-rich regions associated with repeats(*14*).

SVs, such as insertions, deletions, and inversions, contribute disproportionately to human genetic diversity by affecting more genomic sequences than SNVs and can significantly disrupt genes and regulatory elements, leading to relatively larger effects on gene expression and phenotype as defined by association studies(*17, 20*). Indeed, the effect size for this particular class has been estimated to be more than order of magnitude greater than single-nucleotide polymorphisms. In humans, many SVs have been strongly implicated in a variety of diseases, such as neurodevelopmental disorders (e.g., 22p11.2 deletion syndrome, 22q11.2DS)(*20*) and coronary heart diseases (*LPA*)(*21*). Conversely, human-specific SVs have been shown to play important roles in the adaptive evolution of our species, including the adaptations to diet(*22*) and the expansion of human neocortex(*23*). Furthermore, several recent studies provided some of the first evidence for adaptive SV introgression(*12, 14, 16*) and reported novel protein-coding genes with positively selected sites within introgressed regions(*14*). Notably, these studies relied primarily on short-read sequencing data, with limited long-read data available, inadequately capturing the full spectrum of variations, particularly in complex loci, due to the intricate and repetitive nature of many SVs(*17, 19, 21, 24*). Therefore, advancements in data and inference approaches are still required for comprehensive analysis of SV evolution.

Highly accurate long-read sequencing technologies have now made it possible to completely resolve complex repetitive regions in the human genome for the first time, including most SVs(*17, 19, 21, 24, 25*). Recent long-read sequencing efforts from the Human Pangenome Reference Consortium (HPRC)(*21*) and the Human Genome Structural Variation Consortium (HGSVC)(*17*) reveal that over 70% of SVs are inaccessible to short-read sequencing, with many novel SVs located in genes or regions associated with known diseases and complex traits. Here, we set out to systematically build a genome-wide SV introgression map by projecting SVs onto introgressed segments identified from both the HPRC Release I (HPRCr1) panel (n=47) and newly generated long-read phased haplotype assemblies from two PNG individuals, PNG15 and PNG16. We begin with a detailed characterization of SVs and SDs for the new Papuan assemblies compared with the HPRCr1 and the latest T2T-CHM13v1.1 reference assemblies. The addition of Papuan genomes is particularly important because they represent a gap in the current human pangenome (HPRC) as these populations were never included as part of the 1000 Genomes Project (1KG)—the sole source of samples for the first two releases. Papuan genomes were previously reported to harbor the greatest degree of introgression(*4, 5, 26*) and, thus, represent the most likely source to discover introgressed SVs.

Using population genetics approaches and simulations, we first construct a comprehensive SV introgression map and delineate the properties of introgressed segments and SVs across this diverse population panel. In particular, we highlight candidate archaic introgressed centromeres in the Papuan assemblies, providing the first glimpse of centromere evolution in archaic hominins. We also leverage a pangenome graph-based approach to genotype variants in a large cohort of short-read samples and report multiple high-confidence adaptive SV introgression signals among the Papuan people, providing new insights into the role of SV introgression in human evolution and biology.

## RESULTS

### High-quality, haplotype-phased Papuan assemblies

We generated high-coverage sequencing data for a female from Western Lowland Papuan New Guinea (PNG15) and a male from the Solomon Islands (PNG16) using PacBio high-fidelity (>31-fold coverage), Oxford Nanopore Technologies (>44-fold coverage), Arima Hi-C (>40-fold coverage), and Illumina short-read platforms (>18-fold coverage) (**Methods**). A principal component analysis using SNVs from short-read genomes places the two PNG individuals, along with published short-read PNG samples(*27, 28*), separately from the 1KG cohort(*29*) (**Fig. 1A**), confirming the unique genetic diversity of this population group. Using these data, we constructed four nearly complete haplotype-phased assemblies (**Table 1**) ranging from 2.89 to 3.01 billion base pairs using Verkko v1.4.1(*30*), with N50 values of 111–136 million base pairs (scaffolded) or 62.4–74.9 million base pairs (unscaffolded), surpassing both the human reference genome GRCh38 and most of the HPRCr1 assemblies (**Fig. 1B**). The PNG assemblies have an average QV accuracy of 44.5 (an error rate of 3.5×10^-5^per base pair, **Table 1**), with limited collapsed and misassembled sequences (0.18% and 0.03%, respectively) (**Tables S1-S3; Methods**). We estimated that >99.3% of each PNG haploid assembly was completely and accurately assembled, which is highly compatible to the T2T-CHM13v1.1 assembly. Alignment to the T2T-CHM13v1.1 reference revealed 91.3‒95.4% unique alignments, with gaps primarily in acrocentric chromosomes and centromere regions (**Figs. S1-S2**). Overall, these high-quality PNG assemblies provide an important resource for comparative genomics research in humans.

**Fig. 1:**
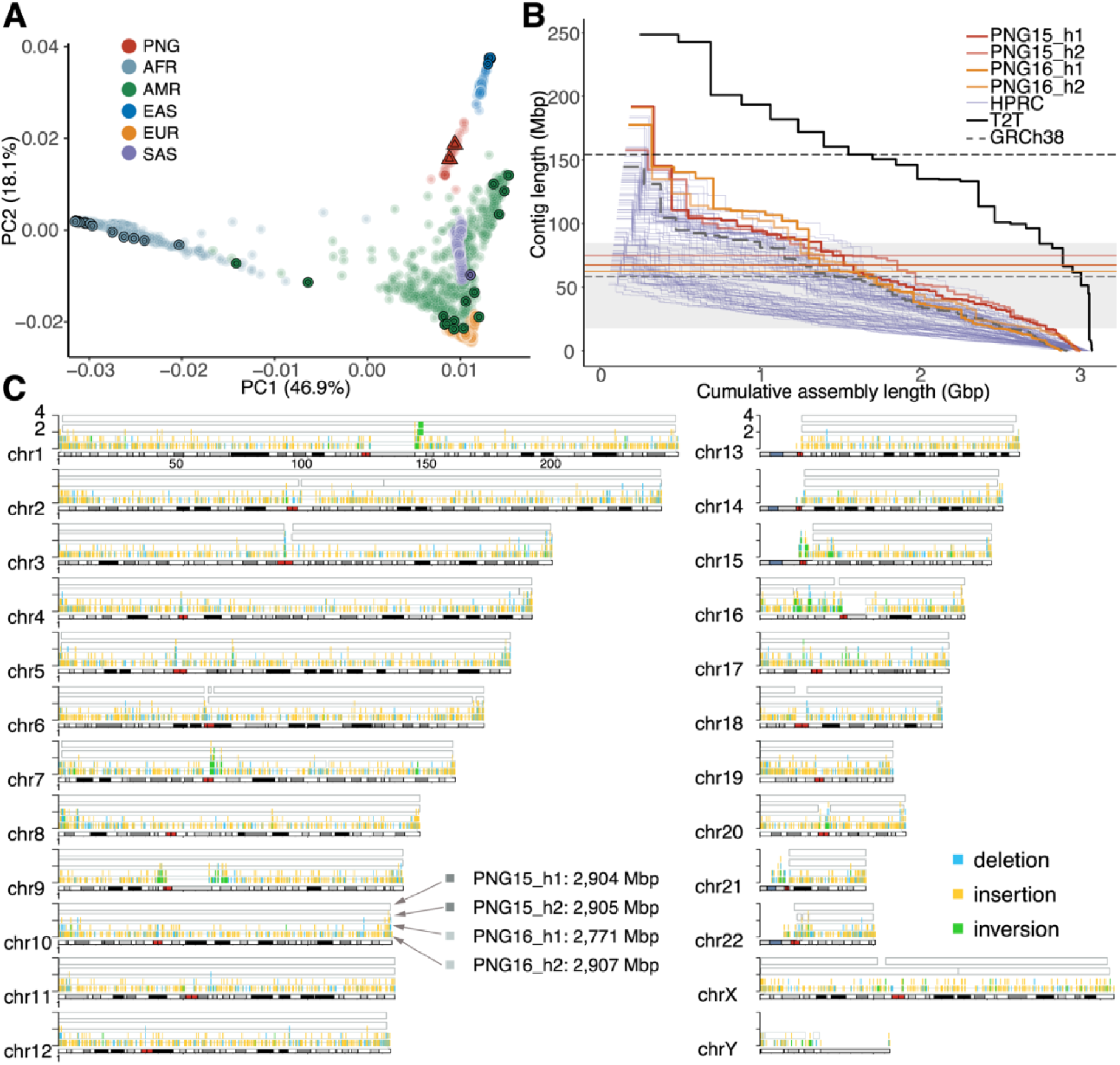
High-quality haplotype-phased diploid assemblies from two PNG individuals. (A) The first two principal components of 71 PNG samples (red), including PNG15 and PNG16 (triangles), along with the 1KG samples. Each point represents a sample, with colors indicating the continent of sampling origin. Bordered symbols indicate HPRC and PNG assemblies. PNG: Papua New Guinea, AFR: African, AMR: admixed American, EAS: East Asian, EUR: European, and SAS: South Asian. (B) Assembly contiguity of the PNG, HPRCr1, GRCh38, and T2T-CHM13v1.1 assemblies. All assemblies are gap (N)-stripped before computation. Red and orange horizontal lines indicate the N50 values for PNG15 and PNG16, respectively. For comparison, black solid and gray dashed lines represent the N50 for T2T-CHM13v1.1 and GRCh38, respectively. The shaded area highlights the range of N50 values for the HPRCr1 assemblies. (C) Completeness of the PNG assemblies is compared with T2T-CHM13v1.1 using contig alignments ≥1 Mbp. The alignment blocks from the top to the bottom are for PNG15_h1, PNG15_h2, PNG16_h1, and PNG16_h2. The histogram within alignment blocks highlights PNG-specific SVs in the assembly discovery set, where the y-axis refers to allele count in the four PNG assemblies and colors indicate SV types.

**Table 1.** Summary of the high-quality haplotype assemblies for two PNG individuals using PacBio HiFi and Oxford Nanopore Technologies (ONT) sequencing. The complete human reference T2T-CHM13 (v1.1) is listed for comparison. The X and Y superscripts indicate the inclusion of sex chromosomes in individual haplotype assemblies.

|  | HiFi read coverage | ONT read coverage | # Contigs (>1 Mbp) | Assembly N50 | Error rate (QV) | % Assembly Error | Gene completeness (BUSCO) | Size of assembly |
| --- | --- | --- | --- | --- | --- | --- | --- | --- |
| PNG15_hap1<sup>X</sup> | 31x | 44x | 108 (40) | 133,306,338 | 3.29x10<sup>-5</sup> (44.8) | 0.44% | 98.77% | 3,005,323,559 |
| PNG15_hap2<sup>X</sup> |  |  | 362 (38) | 111,687,398 | 3.53x10<sup>-5</sup> (44.5) | 0.42% | 98.67% | 3,011,272,601 |
| PNG16_hap1<sup>Y</sup> | 38x | 51x | 146 (48) | 136,876,237 | 3.41x10<sup>-5</sup> (44.6) | 0.51% | 98.19% | 2,894,421,598 |
| PNG16_hap2<sup>X</sup> |  |  | 175 (37) | 129,967,162 | 3.41x10<sup>-5</sup> (44.6) | 0.48% | 98.11% | 3,004,940,457 |
| T2T | - | - | 24 (24) | 150,617,247 | - | - | - | 3,112,236,665 |

### Genomic variation in the Papuan assemblies

We assessed genomic variation in the four PNG haplotype assemblies relative to the human reference genome GRCh38 using both assembly-based callers PAV (v1.1.2)(*17*) and SVIM-ASM (v1.0.3)(*31*) as well as read-based callers Sniffles2 (v2.2)(*32*) and PBSV (v.2.9.0, **Methods**). Because merging variants, especially for SVs, across call sets remains challenging, we chose PAV calls as the primary call set given its high precision and call rates(*17*) and used other call sets as a secondary support. In addition, we included HPRCr1 PAV call set for comparison in our downstream analysis for comparison. For SV comparisons between PNG and HPRCr1 call sets, SVs of the same type that overlap by at least 50% reciprocally are merged(*17*) and re-genotyped based on supporting evidence (**Methods**). PAV identified 4.59 and 4.25 million (or 4.53 and 4.20 million after QC, **Methods**) SNVs and 1.07 and 0.99 million small insertions and deletions <50 bp (indels) in PNG15 and PNG16, respectively, compatible to the number of small variants found in the HPRCr1 non-African samples (mean SNVs: 4.26‒4.40 million, mean indels: 0.97‒1.03 million, **Fig. S3**). Removing variants within the erroneous assembled regions in the PNG assemblies results in ∼1.3% and ∼1.8% decreases in SNVs and small indels, respectively. It is worth noting that compared with the HPRCr1 call set, 12% of the small variants are exclusively found in the two PNG genomes, confirming the presence of unique diversity in the Papuan(*7, 27*).

For large SVs (length ≥50 bp), the suite of callers collectively identified 30,789 and 30,574 insertion (INS), 23,312 and 22,581 deletion (DEL), as well as 175 and 175 inversion (INV) calls in PNG15 and PNG16, respectively. Of the 42,261 insertion and deletion SVs that PAV identified, >89.7% of these SVs (PNG15: 89.7%, PNG16: 90.7%) have at least one secondary support from other callers (**Fig. S4**). Of note, PAV-specific SV calls are significantly longer than those identified by multiple callers (**Fig. S4**, **Table S4**, Mann-Whitney U [MWU] test, p<1.0×10^-7^regardless of SV type)—a likely benefit of assembly versus read-based callers. To assess the SV diversity in the PNG samples, we analyzed PAV SV calls from the PNG and HPRCr1 cohorts together and found that 5.6% of the SVs are PNG-specific (**Fig. 2A**). While the fraction of PNG-specific SVs in the two samples is significantly higher than in any pair of East Asian samples (range: 4.7–4.9%), it is comparable to those observed among admixed American samples (range: 5.2–6.6%). Between SV types, 4% and 6% of the insertions and deletions, respectively, are PNG-specific (**Fig. S5**). We noticed a much lower support rate for inversions (<15%) by different callers and that over 90% of inversions are sample- and/or cohort-specific (PNG vs. HPRCr1), highlighting the difficulty in calling inversions. Because of the unusually high population specificity and the overall lower support for inversions, unless mentioned otherwise, we focus on insertions and deletions for most of the downstream analysis. Similar to previous studies(*17, 21*), we found an inverse relationship between SV length and abundance in the PNG genomes; e.g., 97.6% of SVs are shorter than 10 thousand base pairs (kbp) regardless of SV type (**Fig. S6**). Overall, the PNG-specific insertions and deletions account for 6.8 and 8.6 Mbp, respectively, in the human genome. Among PNG-specific SVs, while 31% (n=1,094 out of 3,568; 12 exon-overlapped SVs) and 32% (n=1,082 out of 3,372; 57 exon-overlapped SVs) of the insertions and deletions overlap with genes (i.e., introns and/or exons), respectively, our analysis shows a significant depletion of these SVs in genes (p<0.001; 1,000 permutations). These results are consistent with the expectation of selection against large SVs in functionally constrained sequences(*17*).

**Fig. 2:**
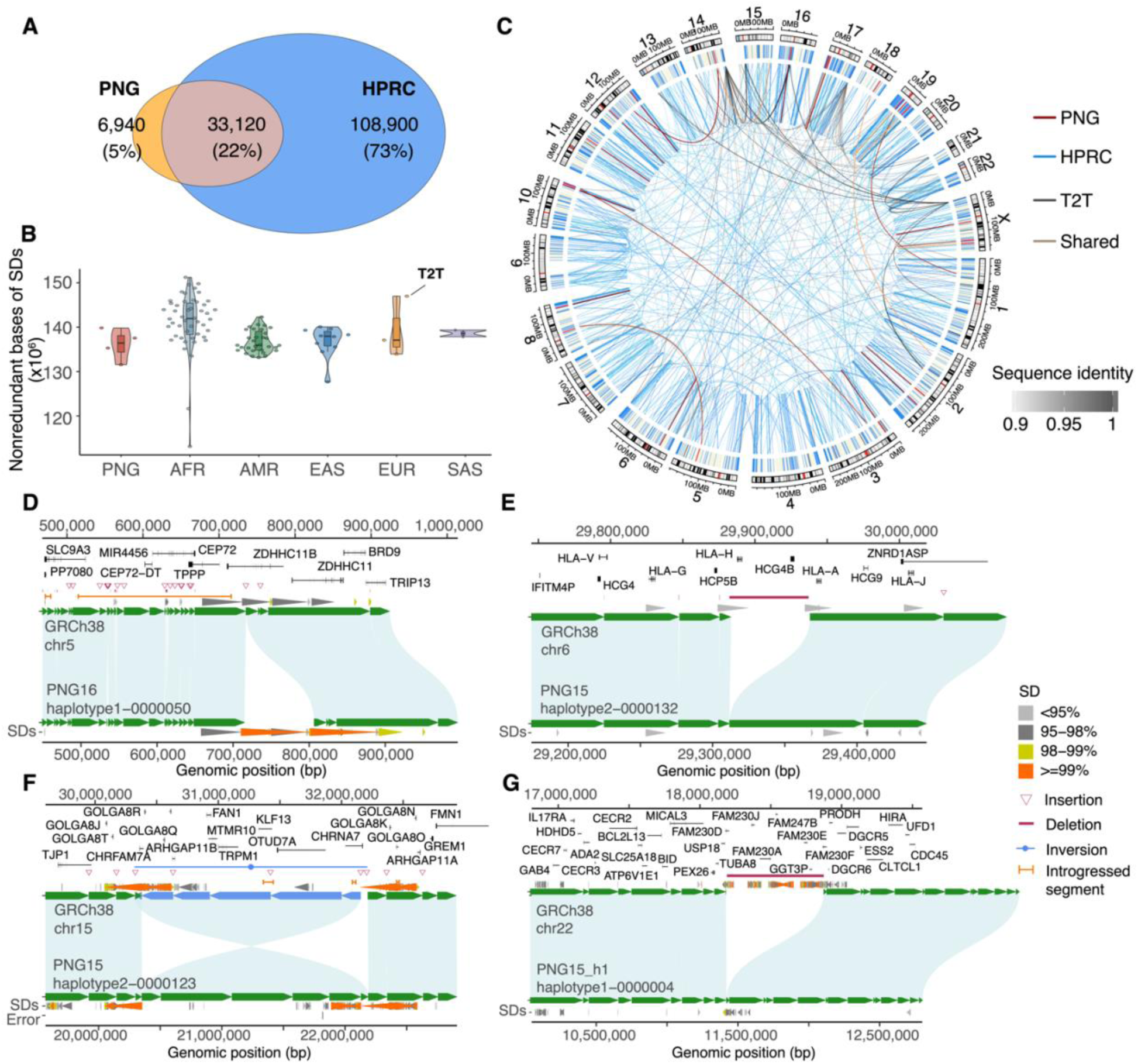
Characterization of genomic variation in the PNG assemblies. (A) Venn-diagram showing the number of SVs shared between PNG and HPRCr1 samples, as well as cohort-specific SV calls. SVs overlapping assembly errors were excluded. (B) Distribution of nonredundant bases of SDs by haplotype across continental groups. SDs are collapsed to compute the total nonredundant bases for each haplotype. The number of such bases is highlighted for T2T-CHM13v1.1 for comparison. (C) Circos plot highlighting intrachromosomal and interchromosomal SDs shared between PNG and HPRCr1 (gray), as well as cohort-specific SDs (red: PNG, blue: HPRCr1). SDs unique to T2T-CHM13v1.1 are black. Color intensity indicates sequence identity between SD pairs. (D-G) Alignment plots between the GRCh38 reference genome (top) and PNG haplotypes showing the cluster of insertions at *TPPP*/*CEP72* (D), the 54 kbp deletion at *HLA-H*/*HCG4B* (E), 1.8 Mbp inversion at chr15q11.3 (F), and 0.68 Mbp deletion at chr22q11.2 (G) loci. The gene annotation of the reference genome is shown at the top, followed by SDs predicted by WGAC (**Methods**).

The high-quality haplotype-phased assemblies also provide a powerful means to assess the diversity and variation of SDs―highly identical sequences (>90% and >1 kbp in length) in humans known to be linked to genomic rearrangements and associated with neurodevelopmental delay and psychiatric disorders(*23, 33*). We generated a genome-wide SD map for the PNG, HPRCr1, and T2T-CHM13v1.1 assemblies and identified 157 Mbp of nonredundant SD sequences (range: 125‒173 Mbp, s.d.: 8.34 Mbp) in the PNG and HPRCr1 assemblies, compared to 182.9 Mbp in T2T-CHM13v1.1. The difference in nonredundant SD bases between T2T-CHM13v1.1 and the PNG/HPRCr1 assemblies is diminished after excluding SDs present within acrocentric/centromeric sequences (**Figs. 2C** and **S7**). At the population level, consistent with previous results(*19*), an excess of nonredundant SD bases is found in African samples relative to others (p=3×10^-9^, one-sided MWU test, **Fig. S7**). No significant differences between PNG and other non-African groups are observed. To determine sample-specific SDs, we conservatively focused on SDs within sequences in assemblies that have ≥1 Mbp of synteny with T2T-CHM13v1.1 and map outside acrocentric/centromeric sequences (**Fig. S8**). Together, we identified a total of 172,624,644 nonredundant SD bases (5.75%) of the human genome. Among these SD sequences, 0.3% (481,154 bp), 0.8% (776,786 bp), and 20.4% (10,639,509 bp) are private to PNG, T2T, and HPRCr1 haplotypes, respectively, while SD bases found in common are at higher frequencies than those private to PNG, T2T, or HPRCr1 haplotypes (**Figs. S9-S13**). PNG-specific SD pairs are significantly higher in identity (one-sided MWU tests, uncorrected p<0.02, except for CHS and YRI) and longer in length (one-sided MWU tests, uncorrected p<0.01, except for YRI) compared with other population-specific SD pairs (**Figs. S14-S15**).

Across all assemblies, 8.2% (n=1,039) of the PNG-specific SVs overlap with SD sequences; 76 of these SVs also affect exonic sequences (**Table S5**). Several notable examples locate at *TPPP*/*CEP72*, *HLA-H/HCG4B*, *TCAF*2, 15q11.3, 22q11.2, *GOLGA8K*/*ULK4P1*, *AMY1A*, *PWRN3*, *CR1*, *GSTM1*, and *APOBEC3A*/*3B* loci (**Figs. 2 and S16-S25**). Among these, in three of the four PNG haplotypes, we observed a cluster of insertions along the *TPPP*/*CEP72* locus (**Figs. 2D and S16**); of which, a 94 bp sequence is inserted in the fourth exon of *TPPP*, which has known associations with diseases such as cystic fibrosis, chronic kidney disease, and multiple sclerosis(*34, 35*). In addition, three of the four PNG haplotypes carry a large, 54,848 bp deletion that removes both *HLA-H* and *HCG4B* (**Figs. 2E and S17**), which play a role in chronic obstructive pulmonary disease and pulmonary function(*36*). Several of these PNG-specific SVs also locate at known large copy number variable loci targeted by natural selection(*22, 37*). At the *TCAF* locus, for example, while structurally similar to a previously published haplotype (e.g., VMRC53_hapB, **Fig. S18**), PNG15 haplotype 1 carries an extra copy of *TCAF2* compared to other known Melanesian *TCAF* haplotypes (e.g., VMRC73_hapA, **Fig. S18**), showing previously uncharacterized haplotype diversity at this locus in Oceania(*22*).

One of the most diverged SD loci is located within a 1.8 Mbp inversion on PNG15 haplotype 2 at chromosome 15q13.3 (**Fig. 2F**), a region known for recurrent inversions(*38*) and microdeletions associated with idiopathic epilepsies(*39*). While no large deletions or duplications are observed, this inversion effectively relocates and reorients a large number of genes, including *ARHGAP11B*, a human-specific gene playing a crucial role in human neocortex expansion(*40*). Another diverged SD locus found in the PNG15 haplotype 1 overlaps with a 0.68 Mbp deletion at chromosome 22q11.2 (**Figs. 2G and S19**), within the low-copy repeat 22A (LCR22A) region. While we found three HPRCr1 haplotypes present similar deletions at this locus with breakpoints off by 10 kbp from each other, further sequence analysis supports that these deletions share an ancestral origin (**Fig. S19**). Our analysis also shows the breakpoints of this deletion locate within homologous exonic sequences of the paralogous genes *FAM230D* (the eighth exon; chr22:18,185,813-18,186,274, GRCh38) and *FAM230F* (the eighth exon; chr22:18873303-18873780), mapping to pair of SDs that share >98.8% sequence identity. We infer this deletion is likely an ancestral state as it presents the same sequence structure as a high-quality chimpanzee assembly (**Fig. S20**)(*41*). This region defines the proximal breakpoint of the well-known morbid copy number variant locus associated with 22q11.2 deletion syndrome (22q11.2DS or DiGeorge syndrome), the most common microdeletion syndrome, affecting ∼1 in 4,000 live births(*20*). Notably, this 0.68 Mbp deletion creates a much shorter SD at LCR22A responsible for driving the majority of 3 Mbp deletions of 22q11.2DS(*20*). Because this deletion essentially removes one of the LCR22 sequences and our sample consists of phenotypically healthy individuals, we hypothesize that this deletion haplotype would significantly reduce the prevalence of the recurrent rearrangement associated with 22q11.2DS among the PNG. These findings highlight the importance of expanding genomics into understudied populations for a better understanding of human genomic variation and show that the PNG haplotype-phased assemblies carry substantial and complex genomic diversity that has not yet been fully characterized.

### A global map of genomic introgression from archaic hominins

Many modern-day non-African genomes are known to carry 1–4% of archaic hominin DNA, e.g., Neanderthal (NDL) and/or Denisovan (DNS), with a higher amount found in Papuans(*5, 7, 27*). To identify SV introgression, we applied two complementary approaches to identify archaic sequences in the modern-day human autosomes and projected SVs onto the inferred introgressed haplotypes (**Methods**). We focused on autosomal data unless mentioned otherwise and first applied Sprime(*5*), a statistic sensitive to archaic sequences carrying unusually higher sequence divergence and extended linkage disequilibrium. Because 24% of the segments found in the PNG samples have a derived allele match rate of <10% to archaic-specific alleles (**Figs. S26- S27**, **Methods**), we conservatively removed these low-affinity segments in the two PNG genomes to minimize potential false positives. The broad range of archaic allele fractions observed across putatively introgressed segments (**Figs. 3A** and **S26-S27**) supports multiple introgression events underpinning these segments in the past(*5*). As a second approach, we apply a hidden Markov model (HMM)(*42*) that leverages existing high-coverage archaic genomes(*1–4*) directly searching for archaic-like sequences in order to further refine putative introgressed segments (**Methods**). In both analyses, we initially focused on segments outside known complex regions, such as acrocentric and high-identity SD sequences (>99% sequence identity), to avoid false positives due to inaccurate mapping (**Methods**).

**Fig. 3:**
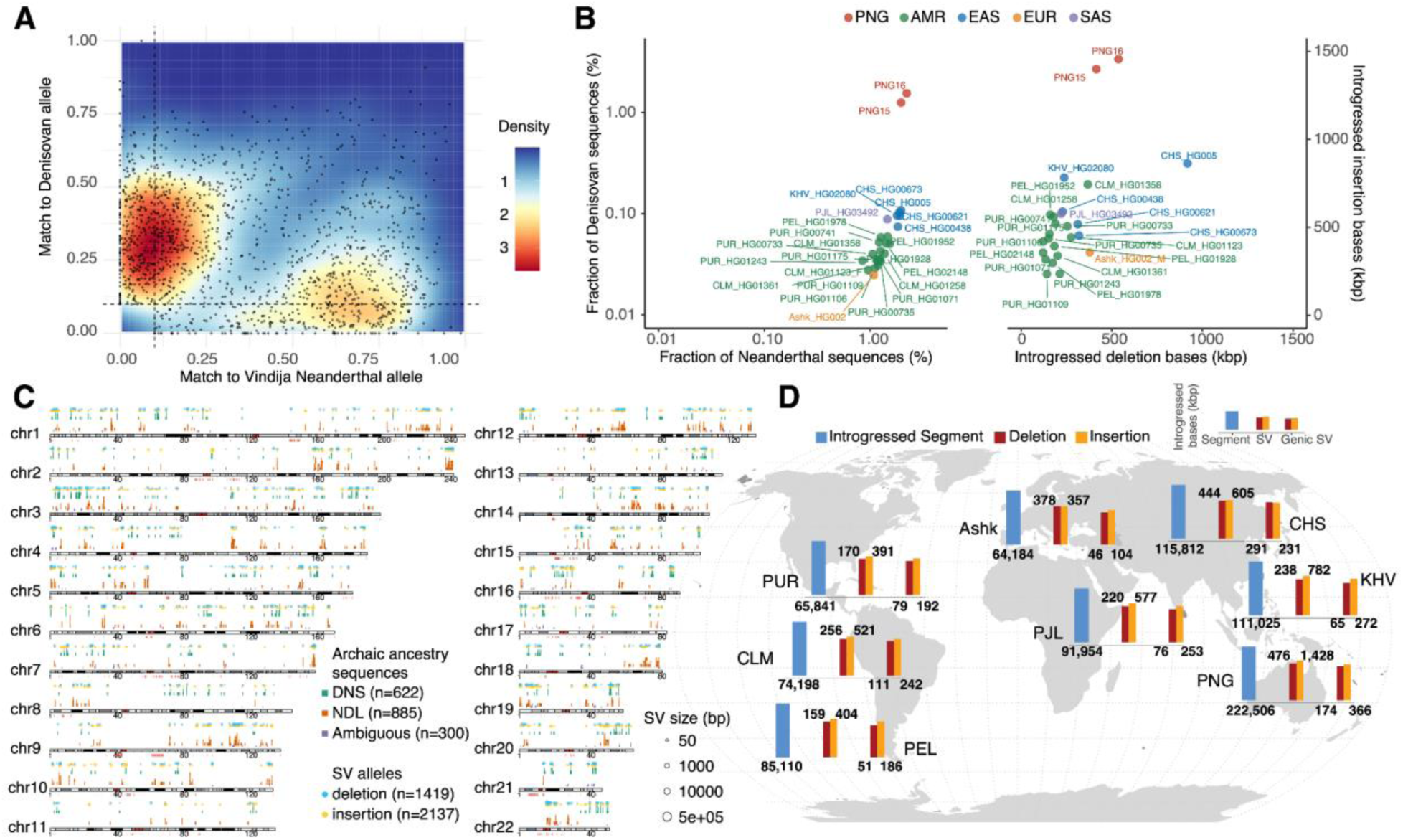
Archaic introgressed sequences and SVs from a diverse population panel of non-African samples in HPRCr1 and PNG cohorts. (A) Density plots of matched proportion to the Vindija Neanderthal and Denisovan genomes in individual introgressed segments (dots). The proportion of putative archaic-specific alleles in each segment was computed for a given archaic genome. Segments with less than 10% match rate on both axes were excluded due to low confidence. (B) Summary of introgressed sequences and SVs in the PNG and HPRCr1 non-African samples. Left: Fractions of Denisovan (y-axis) and Neanderthal (x-axis) sequences in individual diploid genomes. Right: The amounts of introgressed bases for insertion (y-axis) and deletion (x-axis) alleles in individual diploid genomes. (C) Genome-wide distribution of introgressed segments (vertical bars; green: putative DNS sequences, red: NDL sequences, dark blue: ambiguous origin) and SV alleles (circles; blue: deletion, yellow: insertion) in PNG samples. Similar plots for other non-African HPRCr1 samples are in **Figs. S34-S37**. (D) The mean number of bases for introgressed sequences, introgressed SVs, and introgressed SVs that overlap with genes in individual populations. The numbers are expressed in thousands of base pairs. The height of each bar chart is on a base-10 logarithm scale. Ashk: Ashkenazi, CHS: Han Chinese South, CLM: Colombian in Medellin, KHV: Kinh in Ho Chi Minh City, PEL: Peruvian in Lima, PJL: Punjabi in Lahore, PUR: Puerto Rican.

In total, our analysis identified 445 Mbp of introgressed sequences from the four PNG haplotypes, which span over 377 Mbp of the genome. This corresponds to 3.82% and 4.37% of the PNG15 and PNG16 genomes, respectively, predicted to have archaic origins (**Figs. 3B** and **S28**). Specifically, 1.96% and 2.22% of the PNG15 and PNG16 diploid genomes, respectively, likely originate from Neanderthals, with 1.25% and 1.55% derived from Denisovans, with additional 0.6% sequences showing ambiguous archaic ancestry (**Fig. S28**). Segments with ambiguous archaic ancestry are likely due to the actual source populations of archaic admixture and complex demographic dynamics during hominin evolution(*43, 44*). In all cases, the length distribution of introgressed segments is wide with a long tail (median NDL track: 38.8 kbp, s.d.:141 kbp; median DNS track: 33.3 kbp, s.d.: 113 kbp), indicating that these segments are likely resulting from admixture events that occurred at different times in the past (**Fig. S28**)(*5–7*).

Taking advantage of our haplotype-phased assemblies, we constructed a map of introgressed SVs in the PNG samples by physically projecting all SVs identified onto the introgressed haplotypes. Our analysis identified 1,419 deletions and 2,137 insertions mapping to 1,807 putatively introgressed segments (or 176.7 Mbp) potentially derived from Neanderthal- and/or Denisovans-like hominins (**Fig. 3C, Table S5**). While the two PNG genomes carry similar amounts of introgressed insertion bases (∼1.4 Mbp), the number of introgressed deletion bases in PNG16 is 28% more than that in PNG15 (**Fig. S29**). In the PNG samples, twice as many deletions were of Neanderthal origin compared to Denisovan, whereas no such difference was observed for introgressed insertion alleles (**Fig. S29**). In addition, the length distributions of introgressed insertions and deletions were similar between the two PNG genomes (**Fig. S29**).

To build a comprehensive view for the extent of introgressed SVs in modern-day humans, we applied the same analysis to the non-African genome assemblies from the HPRCr1 cohort. Overall, our analysis showed that the PNG samples carry twofold more DNA derived from archaic hominin compared to other non-African HPRCr1 samples (mean fraction: 4.10% in PNG, 1.34% in AMR [admixed American, range: 0.94–1.68%, n=16], 2.13% in EAS [East Asian, range: 2.04–2.19%, n=5], 1.19% in EUR [European, n=1], and 1.72% in SAS [South Asian, n=1]) (**Figs. S30-S33**, **Table S6**). Collectively, ∼335 Mbp of nonredundant introgressed DNA, comprising 11.2% of the genome, were found in these 25 individuals. Stratified by archaic origins, we saw that, on average, while the HPRCr1 individuals carry much less (<0.11%), and on average shorter, putative Denisovan DNA than the PNG (>1.25%), the EUR and SAS individuals carry longer Neanderthal segments than the others (**Figs. 3B** and **S34**). We also identified genome-wide introgressed SVs across population groups (**Figs. S35-S38**). As expected, there is a strong linear relationship between the amount of introgressed SV allele bases and the introgressed fraction among the samples (p=5.3×10^-8^, generalized linear model, chi-squared test, d.f.=1; **Fig. S39**). Across continental groups, on average, introgressed SVs affect 0.6 to 1.9 Mbp per individual, with the highest in PNG (mean: 1,778 SV alleles) and lowest in admixed Americans (mean: 521 SV alleles), while the total introgressed SV bases among individuals are compatible (**Figs. S40-S43**). Across samples, we found that the amount of introgressed insertion bases is notably greater, more than threefold that of deletion bases in some cases (**Figs. S39**). This aligns with the general trend of more insertions than deletions observed in assemblies relative to GRCh38, likely due to reference bias, which favors the discovery of insertion variants over deletions(*45*). Regardless, compared to others, the PNG samples show higher levels of both total introgressed SV bases and the ratio of introgressed insertion bases to deletion bases (**Fig. S39**).

Across all assemblies, 44% of the introgressed SVs (n=1,592) overlap with genes (**Fig. 3D**). It is worth noting that we observed enrichment of introgressed SVs in genes in each continental group, suggesting a potentially functional role of these introgressed SVs. For example, in the PNG samples, 710 introgressed SVs are located within less than 1 kbp of genes (RefSeq release 109, **Table S5**), with an odds ratio of 1.09 (p=0.0056, one-sided Fisher’s exact test; 95% C.I.: 1.04–1.15) for significant genic enrichment of introgressed SVs (p=0.0056, one-sided Fisher’s exact test; 95% C.I.: 1.04–1.15). We also found overrepresentations in multiple gene ontology (GO) biological processes, such as negative chemotaxis (GO:0050919, 4.61-fold enrichment, Bonferroni’s p=2.2×10^-2^, Fisher’s exact test), calcium ion transmembrane transport (GO:0070588, 2.54-fold enrichment Bonferroni’s p=6.6×10^-3^, Fisher’s exact test) and neuron projection guidance (GO:0097485, Bonferroni’s p=3.1×10^-3^, Fisher’s exact test) suggesting potential functional consequence to introgressed SVs (**Table S7**).

We observed several previously reported introgressed SVs, such as a Neanderthal-introgressed compound SV with a 4 kbp deletion and a 31 kbp duplication at the *TNFRSF10D* locus(*14*) and an introgressed variable number tandem repeat (VNTR) haplotype at the *MUC19* locus(*46*) (**Fig. S44**) with an uncertain archaic origin. Of note, we identified multiple novel introgressed SVs; for example, compared to the non-introgressed haplotype, all three Neanderthal-like PNG haplotypes at the *TPPP*/*CEP72* locus encompass the cluster of PNG-specific insertions among the assemblies discussed earlier (**Figs. 2D and S16**). Our sequence analysis suggests that this cluster of insertions represents differences in VNTRs at this locus between modern humans and Neanderthals. We also observed a 52 kbp deletion associated with a Neanderthal-like haplotype, which effectively removes both *GOLGA8K* and *ULK4P1* from the distal site of the chromosome 15q13.3 deletion region (**Fig. S21**). While the syndrome is rarely involved with this distal site(*47*), because *GOLGA* paralogs are known to mediate pathogenic microduplications and deletions at 15q11–13(*33*), this 52 kbp deletion could affect the susceptibility to genomic instability at this locus.

### Adaptive introgression in PNG genomes

The high-quality comprehensive SV introgression map that we generated above makes it possible to further study the evolutionary significance of introgression. We hypothesized that introgressed SVs that rose to high frequencies in the PNG could be the targets of positive selection for local adaptations(*48*). We leveraged the haplotype-resolved PNG and HPRCr1 assemblies, together with PanGenie(*49*) (**Methods**), a pangenome graph-based approach, to construct an augmented pangenome reference, which enables comprehensive genotyping of a broad range of genetic variants, including SVs, in existing large short-read sequencing cohorts from global populations. To maximize the genotyping accuracy and sensitivity, we opted to genotype using a pangenome graph of unmerged variants from the PNG and HPRCr1 assembly call sets, including 28,092,023 SNVs, 8,841,949 short indels, and 613,375 SVs (**Methods**)(*49*). We genotyped these variants in 71 published high-coverage PNG short-read genomes (>20×)(*27, 28*), along with 703 African and 585 East Asian samples from the high-coverage 1KG data set(*29*) and the four high-coverage archaic genomes. Our QC analysis showed an overall 99.1% genome-wide concordance rate for SNVs across samples (range: 98.5–99.3%, **Fig. S45**). Among SVs, the per-sample concordance rates for insertions are highly compatible regardless of size (range: 96.9–98.7%); in contrast, we observed a negative relationship between concordance rate and size for deletions (**Fig. S45**). Across the genome, discordant genotypes tend to occur in subtelomeric and centromeric regions, indicating a limited ability to accurately genotype highly repetitive variants, such as VNTRs (**Fig. S46**)(*17*). Unless specified otherwise, we conservatively excluded discordant variants, those in challenging regions, and those with missing genotypes, leaving 19,222,414 SNVs, 5,345,942 indels, and 199,917 SVs for subsequent analyses.

To detect signals of selection and introgression, we applied various population genetic statistics and controlled for potential biases due to population history in the test results using coalescent simulations based on published demographic models for Papuans (**Methods**). Overall, we identified 54 regions across the genome that show significant signals of positive selection (**Fig. S47**, **Table S8**; length range: 5,134–95,149 bp), of which 18 overlap with exonic sequences and also show evidence of archaic origins (**Table S8**). Variations in several of these genes have been shown to associate with immunity (*CD53*), digestion of lipids (*MALRD1*), neurodevelopment (*NECAB1*), psychiatric disorders (*AKAP11*, *DENND1A*), spermatogenesis (*IQCH*), and stroke (*LRCH1*)(*50–55*). One of the strongest adaptive introgression signals is found at the GBP2/GBP7 (interferon-induced guanylate-binding proteins 2 and 7) locus in the chromosome 1p22.2 region in the PNG (population branch statistic [*PBS*] > 0.75, p < 0.004; introgression statistic *f_D_* > 0.41, p < 0.008). This locus was recently identified as the strongest selection signal in a study using the same PNG short-read genomes(*28*) and likely associated with immunity against diverse pathogens(*56*).

To identify for adaptive introgressed SV candidates, we search for SVs that show significant differentiation in frequency in the PNG cohort compared to the AFR and EAS groups (simulation-based p < 0.05, **Methods**) and located in genomic regions showing evidence for both selection and introgression based on SNV analysis. Across the genome, 13 regions encompassing 16 SVs showed significant selection signals in the PNG cohort (median *PBS_SV_* = 0.79, p < 0.002; genome-wide top 1% *PBS_SV_* cutoff=0.39, **Fig. 4**). All but one of these highly differentiated SVs are shorter than 1 kbp in length (median: 319 bp, range: 52–395,265 bp); 14 of these SVs are insertions and 2 are deletions (**Table S9**). By projecting these selected SVs onto the putatively introgressed loci, we find that two of the selected regions are significantly associated with a Denisovan background, while three are linked to a Neanderthal background (p-value of *f_D_* < 0.034), suggesting that these SVs may have archaic origins.

**Fig. 4:**
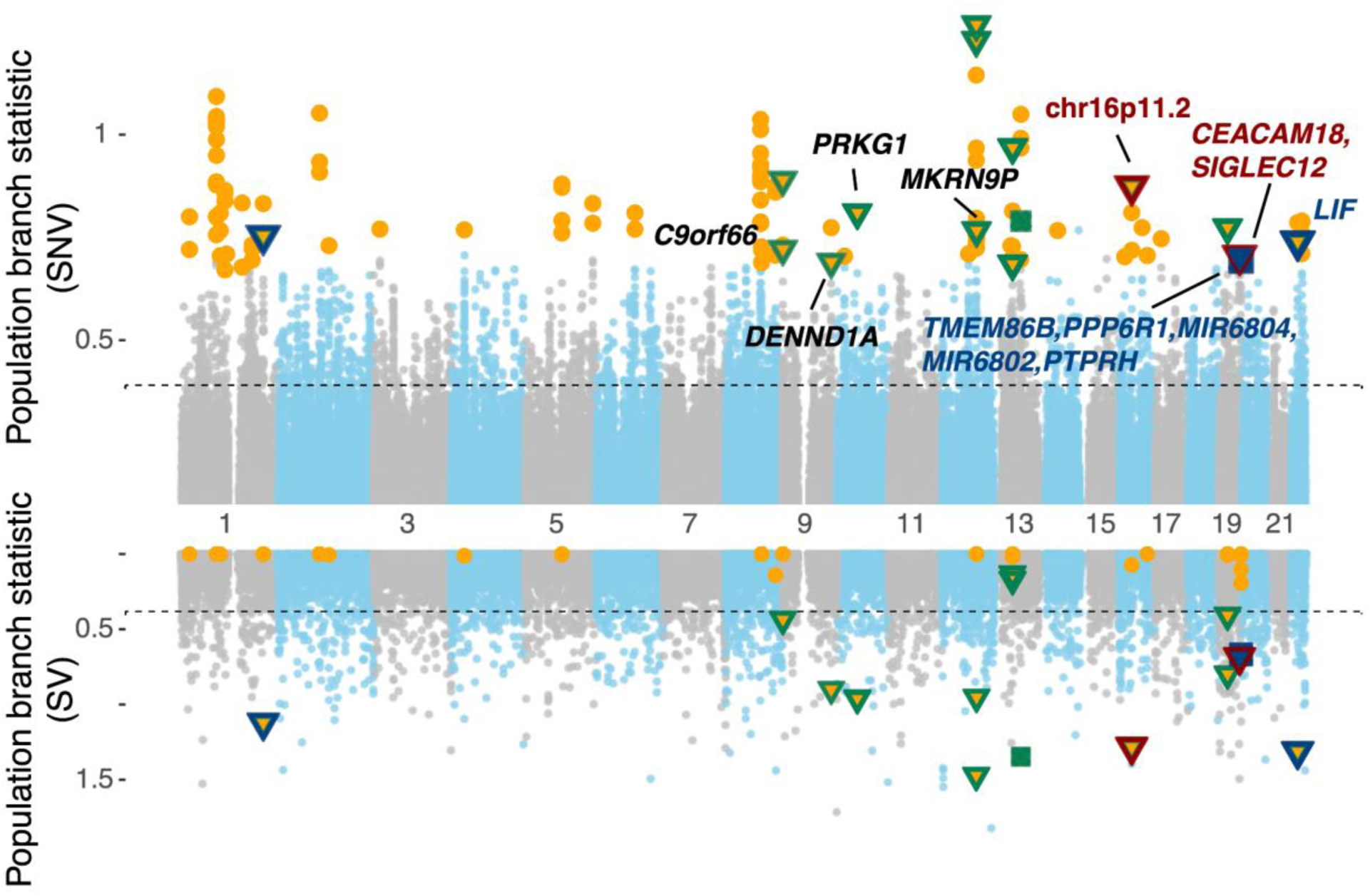
Candidate loci with significant adaptive introgression signals. Top: Genome-wide Manhattan plot (SNVs). Bottom: Hudson (SVs) plot for the adaptive introgression scans in the PNG. The Manhattan plot illustrates the distribution of population branch statistics (PBS) from segments of 100 SNVs, while the Hudson plot depicts the PBS values of individual SVs. PBS values were computed using East Asian and African groups as the sister- and out-groups, respectively. Orange circles represent segments showing significant PBS values (p-value<0.05, simulations based on 10,000 demographic models). Green symbols indicate significant test statistics for SVs (p < 0.05). Red and blue symbols represent putatively introgressed SVs using the Denisovan and Neanderthal as the archaic reference, respectively. Triangles and squares denote insertion and deletion loci, respectively.

Our analysis revealed a putative Denisovan insertion allele (chr19:51484682) at an introgressed locus (chr19:51343160-51528394) on the chromosome 19q13.41 region (**Fig. 5A**). This 61 bp insertion shows an exceptionally higher frequency in the PNG cohort than other groups (68% in the PNG vs. 13% in other groups) and is located in the 4th intron of the gene *CEACAM18* and 6,545 bp away from the transcription stop site of *SIGLEC12*. *CEACAM18* is a member of the *CEACAM* gene family that serves as receptors for bacterial pathogens, such as *Haemophilus* and *Neisseria*(*28*). The absence of other highly differentiated nonsynonymous SNVs around this locus suggests that the insertion allele is a likely target for selection. To further investigate the evolutionary history of this insertion locus, we inferred local genealogies of this locus using pairwise coalescent decoding (**Methods**). Our inference using SNVs in strong linkage to the insertion allele (e.g., chr19:51484850, *r^2^* > 0.96, *D′* = 1) suggests a rather deep divergence (mean: 1.4 million years ago [mya], range:0.99–2.77 mya) from other haplotypes. Pairwise coalescent decoding shows an enrichment of recent TMRCA (time to most recent common ancestors) that are indicative of selective sweeps (**Fig. 5B**). Notably, we found significant evidence for strong selection associated with the insertion-carrying haplotype (s = 0.008, log-likelihood ratio = 6.8468, p = 0.0088, chi-squared test with d.f. = 1). The allele frequency trajectory suggests that the selected variant may have been present at an intermediate frequency (∼0.2) in the population prior to its recent increase (**Fig. 5B**). Consistent with this observation, our analysis of multiple selection episodes indicates stronger support for a selective sweep occurring within the last 100 generations (s = 0.0125, log-likelihood ratio = 10.84, p = 0.0126, chi-squared test with d.f. = 3). *CEACAM18* is exclusively expressed in the small intestine (terminal ileum), and the insertion linked SNVs are also eQTL associated with increasing the expression of this gene (p<9.4×10^-21^, normalized effect size=0.52, GTEx portal) but not associated with the expression of *SIGLEC12*. Given that this gene belongs to the immunoglobulin superfamily, which exhibits evidence of recurrent natural selection on protein surfaces targeted by bacteria in primates(*57*), we hypothesize that the intron insertion allele is likely involved in a pathogen-driven evolutionary process in the PNG population.

**Fig. 5:**
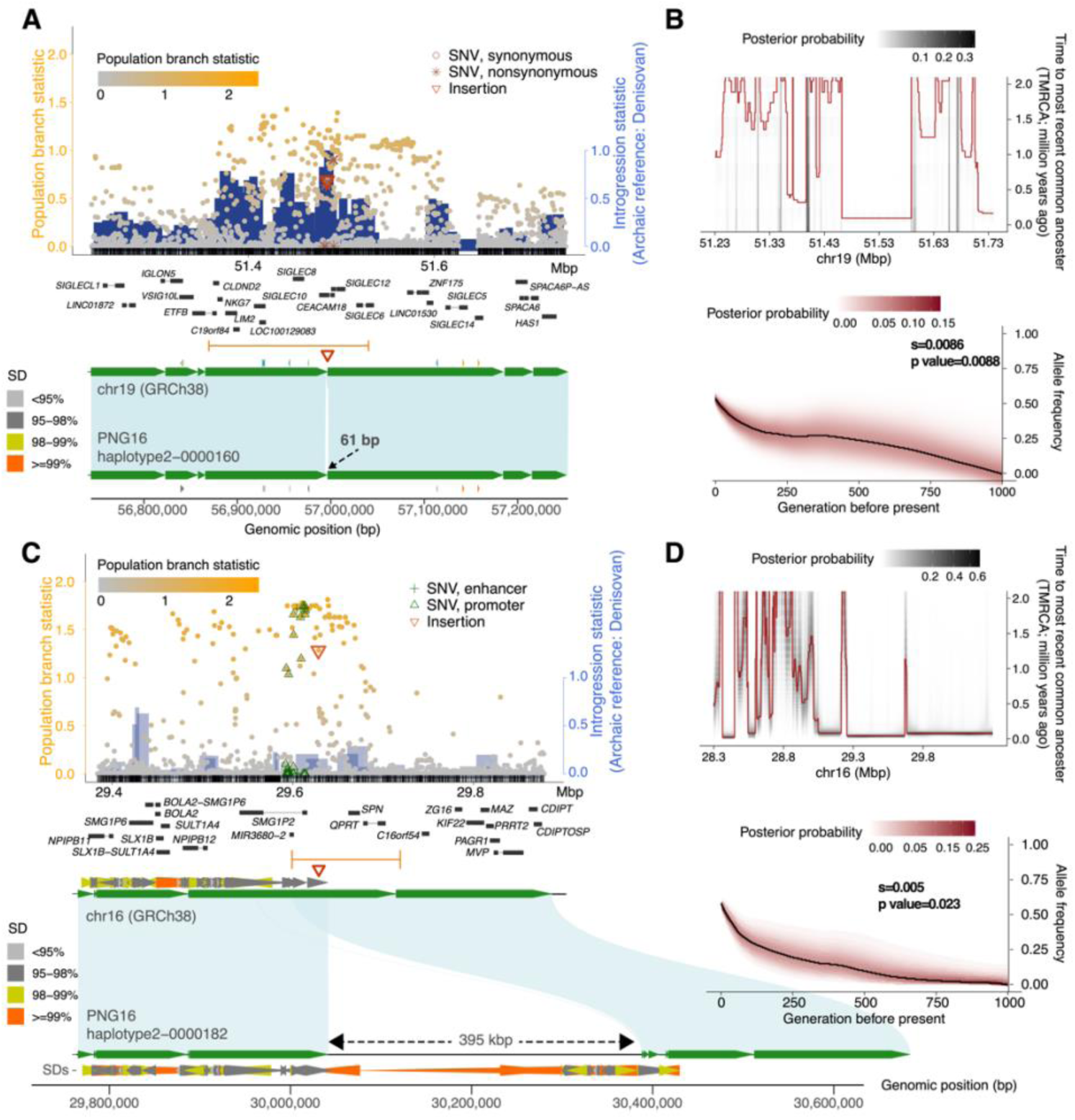
Candidates of adaptive introgressed SVs in the PNG. (A) Distributions of population branch statistics (PBS) of SNVs (circles) across the 61 bp insertion locus at chr19:51231472-51741792. Orange gradient of the circles and the blue histogram indicate the strength of the selection (PBS) and introgression (*f_D_*, archaic reference: Denisovan). Nonsynonymous, synonymous, promoter, and enhancer SNVs as well as SVs are annotated as symbols (RefSeq and ENCODE elements). The middle panel shows gene annotations in the GRCh38 coordinates. Introgressed segments and SVs are denoted as orange segments and red triangles, respectively. Bottom: green ribbon depicts syntenic alignments between GRCh38 and PNG16 haplotype 2 contig, while green arrows indicate sequence orientations. (B) Top: evidence for positive selection using pairwise coalescent decoding between homologous introgressed sequences (**Methods**). The heatmap shows the posterior probability distribution of piecewise time to the most recent common ancestor (TMRCA). An enrichment for recent coalescence times in the distribution of TMRCAs along the sequence is consistent with positively selected loci. Bottom: the likelihood-based ancestral recombination graph approach (CLUES, **Methods**) to infer selection coefficient and allele trajectory using SNVs (e.g., chr19:51484850) that are in linkage with the SV candidate. The heatmap shows the posterior probability of the allele trajectory over the past 1,000 generations. The significance of the selection signal is computed using a chi-squared test (log-likelihood: 5.1641, degree of freedom=1). (C) and (D) are similar to (A) and (B), respectively, but for the putative candidate of the 395 kbp insertion sequence at chr16:29362357-29885651.

Of note, while all three Neanderthal SV candidates are intergenic, a 188 bp insertion is located within 10 kbp upstream of the transcription start site of *PTPRH* (receptor-type tyrosine-protein phosphatase H), a gene recently implicated in regulating intestinal immunity through the dephosphorylation of *CEACAM20*, another member of the *CEACAM* gene family(*58*). Given that the insertion site is within regions displaying strong H3K27ac signals (UCSC Genome Browser, chr19:55218485-55218485), it suggests a potential regulatory role in gene expression. Taken together, our findings support the notion that archaic introgression has contributed to immune adaptations in modern human populations in this region.

The largest introgressed SV found in these PNG assemblies remains a 395 kbp PNG-specific insertion allele at the chromosome 16p11.2 locus (**Fig. 5C**) and was previously reported(*14*). Found in three of the four PNG assemblies (**Fig. S48**), this insertion allele has a frequency of 62.6% in the PNG cohort and is absent in all other samples. We estimated that the insertion haplotype (chr16:29425843-29825843) diverged from the others about 596 thousand years ago (kya; range: 322–870 kya), consistent with its Denisovan origin(*14*). The fully phased haplotypes allow us to accurately decode pairs of first coalescence among haplotypes to estimate the local states of time to most recent common ancestor (TMRCA) (**Methods**). Between the insertion haplotypes, we observed low TMRCA estimates (<120 kya, **Fig. 5D**) around the insertion site (chr16:29628407) compared with the flanking sequences but did not find such signals in other populations (**Fig. S49**). The estimated low TMRCA among insertion-carrying haplotypes aligns with the timing of the inferred introgression event (60–170 kya)(*14*) and is indicative of a clear pattern of a selective sweep—a recent burst of coalescence among the derived lineages(*59*). This insertion encompasses the gene *NPIPB16*, where multiple amino acid substitutions are likely targets of selection(*14*).

Here, we leveraged our population sample and the likelihood-based method CLUES(*60*) to infer the time and strength of selection (**Methods**). Using SNVs in strong linkage to the insertion allele (e.g., chr16:29625843 and chr16:29626149, *r^2^* > 0.41, *D′* > 0.86), we found significant evidence for strong selection (s = 0.005, log-likelihood ratio = 5.16, p = 0.023, chi-squared test with d.f. = 1, **Fig. 5D**). The inferred allele trajectory suggests that the selected introgressed variant segregated at low frequencies (<0.05) for most of its time in the population and only raised to high frequencies starting ∼200 generations ago. In addition, we explicitly tested a model of multiple selection events and found stronger evidence of selection with a selection coefficient of 0.016 starting at 200 generations ago and changed to 0.005 <100 generations ago (log-likelihood ratio = 18.47, p = 0.0003, chi-squared test with d.f. = 3). Our inference suggests that this insertion allele has remained nearly neutral (s < 0.0028) and at low frequencies for an extended period since its introgression into the PNG population until strong selection began approximately 5,800 years ago (**Fig. 5D**), coinciding with the emergence of agriculture in the region during the mid-Holocene(*61*). While the phenotypic outcome of this selection is unclear, the *NPIP* gene family has been hypothesized in the involvement in innate antiviral responses(*62*) and/or related to other immune- or autoimmune-related functions with evidence of ancient and ongoing positive selection in the human lineage(*63*).

### Potential centromere introgression from archaic hominins in the PNG genomes

The high-quality PNG haplotype-assemblies provide a unique opportunity to directly test the hypothesis of centromere introgression from archaic hominins(*64*). We first aligned our high-quality haplotype-resolved PNG assemblies against the complete reference assembly T2T-CHM13v1.1 to identify PNG centromere sequences. To ensure the quality of these centromeres, we removed contigs that contain gaps (Ns) in sequence, do not traverse into unique sequences, or have large-scale misassemblies as described earlier (**Methods**); 59.8% of centromeres (55/92) of PNG15 and PNG16 are considered completely assembled after quality control (**Table S10**). Using the length of active α-satellite higher-order repeat (HOR) arrays (**Methods**), similar to previous studies(*18*) we found large variation in centromere size across different chromosomes, ranging from 0.98 to 6.20 Mbp (s.d.: 1.34 Mbp, **Figs. S50-S73**, **Table S10**); however, no significant correlations between assembled centromere and chromosome sizes are observed (**Fig. S50**). It is worth noting that some chromosomes, such as 8, 9, 11, 14, and X, are similar in size, while others, such as 7, 10, 19, 21, and 22, show greater difference in size among the PNG assemblies (**Fig. S50**). In contrast, the variability in the size and location of the putative centromeric hypomethylated loci associated with kinetochore attachment across centromeres and samples is relatively limited (**Figs. S51-S73**).

Because extensive centromeric repeat variation limits the accuracy of sequence alignments and variant calling, we developed an alignment-free, k-mer approach to search for evidence of introgressed centromeres (**Methods**). Briefly, leveraging the limited presence of archaic DNA in modern-day Africans, we used the high-coverage Neanderthal and Denisovan genomes and identified archaic hominin (ARC)-specific k-mers that are not present in modern-day African samples. Our method subsequently recorded the count of ARC-specific k-mers within 2 kbp windows across each individual haplotype assembly. To show that these ARC-specific k-mers are informative for introgression signals, we examined if the putatively introgressed segments reported above are enriched with ARC-specific k-mers. In both PNG samples, windows that overlapped with putatively introgressed segments have significantly higher counts in ARC-specific k-mers than those nonoverlapping ones (p<2.2×10^-16^, Mann-Whitney U test), showing that our method is able to detect archaic sequences in the assemblies. Among the 55 complete centromeres, 5 and 6 centromeres (mapping to chromosomes 4 [n=1], 5 [n=1], 9 [n=1], 11 [n=3], 17 [n=2], and 22 [n=3]) show evidence for archaic origins in PNG15 and PNG16, respectively (**Fig. 6**). The strongest signal for centromere introgression occurs on chromosome 4 of PNG16 haplotype 2 (**Fig. 6A**) and chromosome 22 of PNG15 haplotype 2 (**Fig. 6B**) as well as PNG16 haplotypes 1 and 2 (**Fig. S72**). Not only do the ARC-specific k-mer counts exceed that of the genome-wide introgression proportion in PNG16 (4.37%, ARC-specific k-mer threshold = 42), large putatively introgressed segments based on the SNV-based inference are found at both sides of the centromere despite limited availability of unique sequence in the region.

**Fig. 6:**
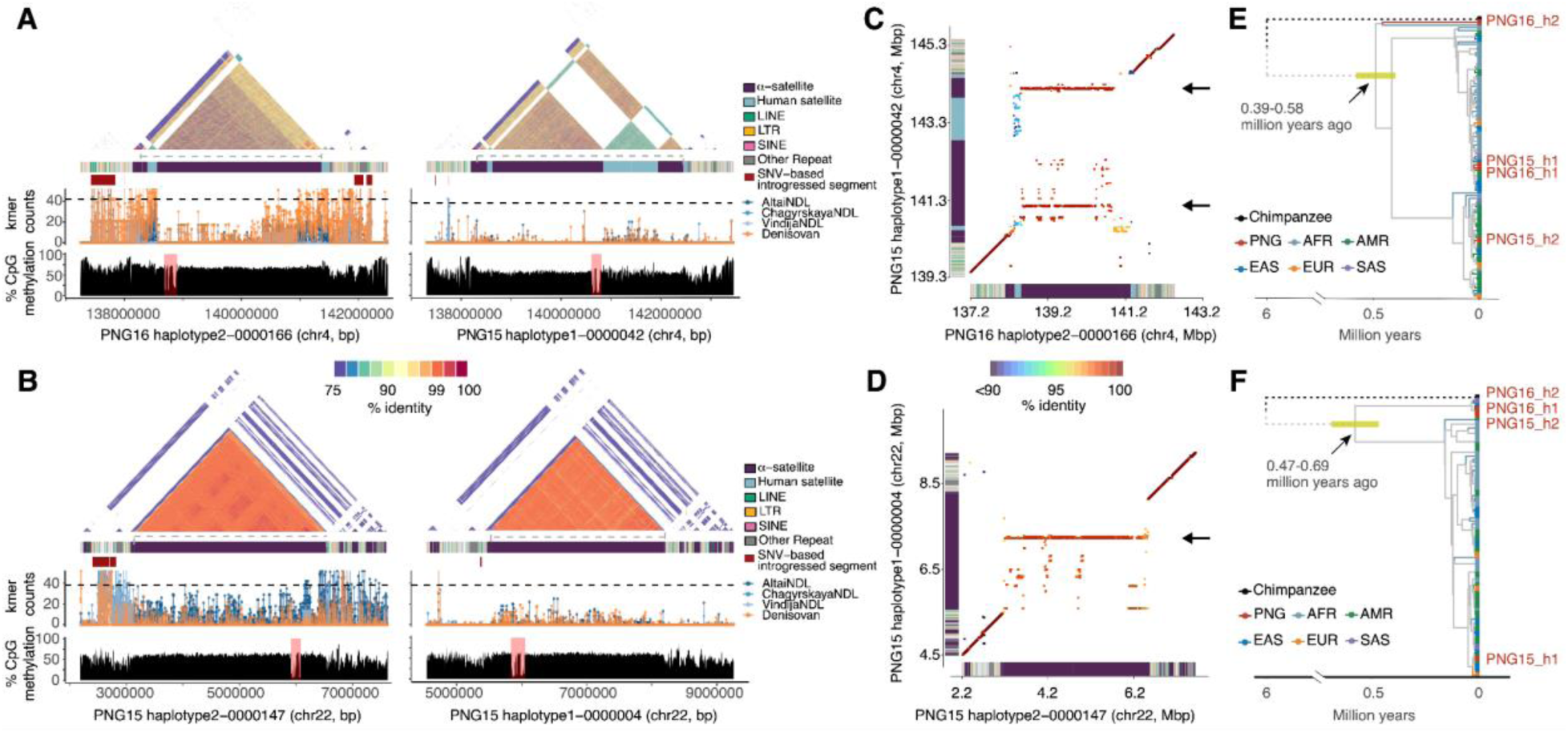
Evidence for archaic introgressed centromeres on chromosomes 4 and 22. (A&B) Comparison between a putative introgressed (left) and non-introgressed (right) centromeres. Top panels: heatmaps of pairwise sequence identity in 5 kbp windows (StainedGlass) across the centromere and flanking regions. Colors of the heatmap indicate sequence identity. The colored horizontal bars show underlying repeat content across the centromere regions. The gray dashed lines indicate the inferred centromere sequence based on annotated active α-satellite arrays. Red rectangles represent SNV-based introgression signals. Middle panels: Distribution of archaic-specific k-mers across 20 kbp windows using four high-quality archaic short-read genomes. The black dashed lines indicate the genome-wide k-mer count cutoffs, 38 and 42, that correspond to the genome-wide archaic introgression fraction for PNG15 and PNG16, respectively. Bottom panels: The percentage of average methylated CpG frequency across 5 kbp bins across the region. The pink rectangles indicate the putative centromeric hypomethylated loci associated with kinetochore attachment. (C&D) Allelic variation between the putative archaic and modern human centromeric haplotypes. 20 kbp windows with a 1 kbp sliding step from the query sequence (horizontal axis) were aligned to the target sequence (vertical axis). Only the best alignments, based on sequence identity, were retained. Colors of dots indicate percent sequence identity. The α-satellite and other human satellite array structures are shown on the axes as in (A&B). The black arrows indicate the modern human α-satellite HORs expanded in the putative archaic centromeric sequences. (E&F) The phylogeny of the putative introgressed sequence from the left flanking of centromere in (A). The ancestral branches leading to the chimpanzee and ancestral lineages of humans are truncated (dashed lines) for illustrative purposes. 95% high posterior density intervals for node ages between 0.2 and 1.0 million years are displayed as horizontal green line

To investigate structural differences between putative archaic and modern human centromere sequences, we performed window-based alignments between archaic and modern human centromeres, archaic centromeres, and modern centromeres (**Figs. 6C&D** and **S74-S75**). While centromeres with the same ancestry origin show relatively higher sequence identity as expected under identity by descent (**Figs. S74-S75**, left panels), centromeres with different ancestry origins substantially diverged in both sequence identity and structure (**Figs. 6C&D** and **Figs. S74-S75**, right panels). Considering both nucleotide and indel differences, in the α-satellite HORs of chromosome 4, the sequence identity between two modern human sequences (median: 99.8%) is significantly higher than that between putative archaic and modern human sequences (median: 98.8%) (one-sided Mann-Whitney U test, p < 2.2×10^-16^). Similarly, in the α-satellite HORs of chromosome 22, the median sequence identity between two putative archaic centromeric sequences is 99.5%, compared to 98.5% between putative archaic and modern human sequences. In addition, the putative archaic α-satellite HORs on chromosome 4 seem to be expanded from two short α-satellite HOR regions in the modern human counterpart (PNG15 haplotype 1-0000042:138,987,977–139,068,718 and 141,985,570–142,102,299, **Fig. 6C**). The putative archaic α-satellite HORs on chromosome 22 also primarily share homology with a narrow region in the modern human haplotype (PNG15 haplotype 2-0000147:7,206,683-7,344,983, **Fig. 6D**). Notably, unlike modern human chromosome 4 centromeric sequences, the archaic haplotype lacks the unique, long stretch of human satellite sequences (**Figs. 6A** and **S54**).

To further support the hypothesis of archaic introgressed centromeric sequences in the PNG samples, we constructed phylogenetic trees using sequences orthologous to the flanking introgressed region from all haplotype-resolved assemblies. Our phylogenetic reconstruction shows that the PNG16 haplotype 2 chromosome 4 centromere lineage diverged from the rest of the modern human lineages 489,447–507,131 years ago (95% highest posterior density [HPD] intervals: 397–583 kya for left flanking and 386–633 kya for right flanking, **Figs. 6E** and **S76**). In addition, we also observed evidence for archaic origins for the chromosome 22 centromeres of PNG15 haplotype 2, PNG16 haplotype 1, and PNG16 haplotype 2 (**Figs. 6F** and **S72**). We noted that in each of these chromosome 22 centromeres only one of the flanking sequences shows introgression signals based on the SNV-based analysis (**Fig. S72**). The lineage that gave rise to the three PNG haplotypes diverged from other modern human lineages approximately 580,661 years ago (95% HPD interval: 469–694 kya, **Fig. 6F**). These divergence time estimates are all consistent with the 400,000–700,000 years of modern human–Neanderthal/Denisovan divergence(*2, 65*), suggesting that these centromeres are likely introgressed from archaic hominins into the early ancestors of the PNG.

Estimating the mutation rate within α-satellite HORs has been challenging(*18*), but the putatively introgressed archaic centromeric sequences offer a valuable recent time point for such estimates. We attempted to use the expansion of modern human α-satellite HORs in the putative archaic centromeric sequences (**Fig. 6C&D**) and assumed orthologous sequences between archaic and modern humans within these expanded loci. To estimate mutation rates, we applied an evolutionary model(*66*), assuming a human-archaic divergence of 400-700 kya based on our phylogenetic inferences (**Methods**). Using this approach, the mean mutation rates of α-satellite HORs for chromosomes 4 and 22 are 2.94×10^-8^ (s.d.: 0.91×10^-8^) and 7.46×10^-8^ (s.d.: 1.89×10^-8^) per allele per generation, respectively (**Fig. S77**). Compared to the mutation rate for the euchromatic portion of the human genome(*67, 68*), our estimated mutation rate for α-satellite HORs is up to 5.96-fold higher, reflecting a 35.4% increase from recent estimates(*18, 69*). While sequence alignments for α-satellite HORs are expected to be biologically meaningful between closely related hominins, we note that these estimates are only approximations, as our approach does not account for the observations of rapid turnover and structural changes in human α-satellite HORs(*18*). Nevertheless, our discovery of putatively archaic introgressed centromeres in the modern human gene pool, along with direct measurements of α-satellite HORs mutation rates, highlights the complexity of centromere and genome biology and opens new avenues for research in human evolution and genome function.

## DISCUSSION

DNA introgression from closely related species is a common biological phenomenon, contributing genetic novelty and facilitating local adaptations in recipient populations. Genomic studies have revealed interbreeding between modern humans and other hominins, such as Neanderthals and Denisovans, with evidence of adaptive introgression at both SNV and SV levels(*10–16, 28, 42, 43, 48*). While the importance of SVs in phenotypic variation and human evolution has long been recognized, studying them has been challenging due to their complexity and the limitations of short-read sequencing(*12, 14, 16, 17, 19, 21–24, 37*). Over 70% of SVs and small variants within repetitive sequences remain undetectable by short-read data, leaving a gap in our understanding of introgressed variations(*17*). In this study, we overcame this challenge by integrating long-read haplotype assemblies from two PNG individuals with high-quality assemblies from 47 HPRCr1 individuals. This allowed us to create a global map of introgressed sequences from Neanderthals and Denisovans, revealing a wealth of previously untapped genomic variation, especially SVs. Our findings raise new questions about the evolution of this variation and provide a framework for further study on their roles in human biology and evolution.

The four newly assembled PNG genomes are more complete than the HPRCr1 assemblies, revealing 12% of small variants and 5% of SVs exclusive to these individuals. These PNG-specific variants affect over 12 Mbp of the genome, three times the number of SNVs found in a typical non-African genome. While the PNG-specific variants are compatible to those from other populations in number, they have several unique signatures, including an enrichment of PNG-specific deletions as well as higher sequence similarity and longer in length for PNG-specific SDs compared to those in other populations although the basis for these differences is unknown. As expected, SVs are found significantly depleted in genic sequences(*14*); yet, the observation of rare SVs at biomedically relevant loci, such as the rare 0.68 Mbp deletion at the 22q11.2 deletion syndrome locus and large 1.8 Mbp inversion at the chromosome 15q13.3 idiopathic epilepsy-associated deletion locus(*39*), calls for the inclusion of more diverse samples in genomics. In the case of the 22q11.2 deletion, for example, with major clinical characteristics of learning disabilities and congenital heart malformations, approximately 90% of individuals affected with the syndrome have a similarly sized *de novo*, ∼3 Mbp hemizygous deletion at 22q11.2(*20*) resulting from meiotic non-allelic homologous recombination events between multiple flanking low-copy repeats termed LCR22A-D. Located at the proximal portion of the typical 22q11.2DS locus, this 0.68 Mbp PNG deletion removes a large portion of the LCR22A locus that encompasses four protein-coding genes (*GGTLC3*, *TMEM191B*, *RIMBP3*, and *FAM246B*) and a dozen RNA genes, including the two right at the breakpoints of this deletion (*FAM230D* and *FAM230F*), which are near multiple DCGR genes (*DGCR2*, *DGCR5*, and *DGCR6*) critical for 22q11.2DS. Although the phenotypic impact of this large PNG deletion remains unclear, given that our samples were collected from individuals who were healthy at the time, the removal of LCR22A may confer protection against genomic instability in deletion carriers by reducing the risk of LCR-mediated unequal crossing-over events. This thus underscores the importance of including diverse populations in genomic research to uncover new variants with potential implications for disease and evolution.

While the present study only included 49 world-wide samples, our global introgression map reveals complex forms of genetic variation of archaic hominin origin that a conventional short-read-based population genetics analysis would miss. On average, introgressed SV alleles affect 0.6 to 1.9 Mbp per individual, with the highest amounts in PNG populations and lowest in admixed American groups. Unexpectedly, PNG individuals carry more Neanderthal sequences than Denisovan, contradicting the expected genome-wide estimates of ∼4% Denisovan and ∼2% Neanderthal ancestry(*5, 7, 27*). This disparity may result from the differences between the sequenced Denisovan genome and the actual Denisovan population that interbred with the ancestors of the PNG as well as complex demographic histories in Oceania(*43*). Notably, our PNG samples are lowlanders, who are known to carry less Denisovan DNA than highlanders(*43*), suggesting that a larger cohort, including highlanders and samples from other regions, would provide a more comprehensive view of SV introgression to better assess the genomic contributions of extinct relatives to the modern human gene pool.

Our analysis of adaptive SV introgression continues the trend of identifying archaic variants contributing to human immunity(*11, 13, 28, 43, 48*). Our top Denisovan-introgressed SV, a 61 bp insertion, is involved in an immune-related gene *CEACAM18*. Our population genetics inferences suggested a 0.5% increase in reproductive success for both SVs. In addition, the 188 bp Neanderthal-introgressed intergenic insertion distal to *PTPRH* could impact the function of the gene in intestinal immunity(*58*). Notably, one of our non-SV adaptive introgressed signals is within the cluster of guanylate-binding proteins (GBP) family, playing a crucial role in immune response mechanisms(*43*) and being previously identified as a candidate for positive selection in the PNG(*28*). This further supports the idea that introgression from archaic populations, particularly Denisovans, has contributed to the immune adaptation of modern human populations in the region. Our recent estimates for the onset of selection of these variants in the mid-Holocene align with the emergence of agriculture in the region(*61, 70*). While agriculture and zoonotic diseases have been suggested as drivers of adaptation(*13*), it is likely that the intensification of cultural practices starting around 5,000 years ago, including the domestication of animals associated with agriculture and regional population movements within the PNG and with other groups in the Wallacean Islands (e.g., East Indonesia) and the Bismarck Archipelago(*70*), played a more significant role in pathogen transmission and genetic adaptation than the emergence of agriculture alone. In conclusion, the introgression of Denisovan and Neanderthal alleles has significantly influenced adaptive traits, particularly in immune response and environmental adaptation. One of the major limitations of our study is the inclusion of only 49 individuals (98 haplotype assemblies), which represents only a small proportion of potential human genetic diversity. Further research with a larger data set and into the functional consequences of these introgressed alleles, especially in centromeric regions and immune genes, will deepen our understanding of how archaic human populations contributed to the survival and adaptation of modern humans in diverse environments.

Consistent with previous studies(*18, 64*), we observed large centromeric variation in length and structure of α-satellite HORs as well as putative kinetochore attachment sites. Given the crucial role of the centromere function in chromosomal stability and integrity during cell division(*71, 72*), our finding of introgressed centromeres in the PNG raises an intriguing question about human evolution. Following interbreeding, introgressed Neanderthal/Denisovan alleles, along with linked neutral alleles, are hypothesized to have been subjected to more efficient purifying selection in the larger modern human population(*8*), especially for functionally conserved regions in the genome(*9*). It remains unclear to what extent these variations contribute to centromere biology and function, and how an archaic centromere has persisted in modern humans to this day. It is likely that the diverse α-satellite HORs among haplotypes suppressed recombination, reducing the effectiveness of purifying selection due to a reduction of local effective population size. Our results suggest that areas of suppressed recombination, including centromeres and inversions, will be regions prone to capture larger swaths of ancient/archaic DNA with much deeper coalescences than most of the genome (e.g., the chromosome 17q21.31 inversion(*73, 74*)), providing unique glimpses into the structure and organization of ancient genomes. Our results demonstrate the potential to recover archaic centromeres by phased/genome assembly of diverse humans providing the potential to compare the structure and function of diverse centromeres and their role in human adaptations.

## Materials and Methods

### Summary

Detailed descriptions for Materials and Methods can be found in Supplementary Information. Briefly, two human lymphoblastoid cell line (LCL) samples (PNG15 and PNG16) were collected in Papua New Guinea with informed consent and ethics approval from the PNG Medical Research Advisory Committee (MRAC 16.21), the University of Melbourne (1851585.1), and French Ethics Committees. A material transfer agreement enabled sharing between the University of Melbourne and the University of Washington. We generated PacBio HiFi, UL-ONT, Illumina, and Arima Hi-C sequencing data for both PNG LCLs. Fully phased assemblies were produced using Verkko(*30*) (v1.4.1), with error annotations using Flagger(*21*) and NucFreq(*75*). Assembly quality and gene completeness were evaluated using Merqury(*76*) and compleasm(*77*). SVs were called using PAV(*17*), SVIM-asm(*31*), PBSV, and Sniffles2(*32*). SDs were annotated using whole-genome assembly comparison(*78*). Archaic introgression sequences of the PNG, along with the HPRCr1(*21*) assemblies, were detected using an HMM-based(*79*) and a reference-free (SPrime(*5*)) methods and refined based on consensus. Additional validation was performed with a k-mer–based method using archaic-specific k-mers. SV genotyping using a pangenome graph of 98 assemblies was performed with PanGenie(*49*) for 1,592 short-read genomes(*1–4, 11, 27, 29, 43*). Population genetics statistics including population branch statistics(*80*), nucleotide diversity(*81*), *f_D_*(*82*), *F_ST_*(*83*), *r^2^*, and Lewontin’s *D’*(*84*) were used for selection analysis, and coalescent simulations with msprime(*85*) were used for determining statistical significance. Pairwise coalescent decoding was used for inferring local genealogies, and selection coefficients were estimated with CLUES(*60*). Centromere structures were analyzed using StainedGlass(*86*). Centromeric methylation profiles were inferred from ONT reads. Phylogenetic dating was performed using BEAST(*87*). Mutation rates were estimated using Kimura’s model of neutral evolution(*88*) with sequence divergence estimated across 20 kbp centromere windows, incorporating a range of divergence times and ancestral population sizes. Full data are available via the European Genome-Phenome Archive (EGASXXXXXX, accession numbers pending approval).

## Acknowledgements

The authors thank T. Brown for assistance in editing this manuscript. Funding: This work was supported, in part, by the US National Institutes of Health (NIH) grant R01HG002385 to E.E.E. P.H. is supported by an NIH Pathway to Independence Award (NHGRI, 5R00HG011041). F.-X.R. and N.B. are supported by the French National Research Agency (ANR) grant ANR-20CE12-0003-01. I.G.R. is supported by Australian Research Council Discovery Project DP200101552. E.E.E. is an investigator of the Howard Hughes Medical Institute.

This article is subject to HHMI’s Open Access to Publications policy. HHMI lab heads have previously granted a nonexclusive CC BY 4.0 license to the public and a sublicensable license to HHMI in their research articles. Pursuant to those licenses, the author-accepted manuscript of this article can be made freely available under a CC BY 4.0 license immediately upon publication.

## Author contributions

P.H. and E.E.E. designed and planned experiments. C.K., F.-X.R., I.G.R., M.L., M.P.C., and N.B. collected PNG samples and constructed cell lines. C.B., K.H., and K.M.M. prepared libraries and generated and analyzed sequencing data. P.H., N.S., D.S.G., A.J., and W.T.H. performed variant calling and bioinformatics analyses. P.H., D.S.G., A.J., N.S., W.T.H., and D.P. analyzed long-read sequencing data and assembled contigs. P.H. designed and performed population genetics inferences and analyses. P.H. and E.E.E. wrote the manuscript with input from all authors.

## Supplementary Materials

Materials and Methods

Supplementary Text

Figs. S1 to S77

Tables S1 to S10

References (89–103)

## References

1. F. Mafessoni et al., A high-coverage Neandertal genome from Chagyrskaya Cave. Proc Natl Acad Sci U S A 117, 15132–15136 (2020).

2. K. Prufer et al., A high-coverage Neandertal genome from Vindija Cave in Croatia. Science 358, 655–658 (2017).

3. K. Prufer et al., The complete genome sequence of a Neanderthal from the Altai Mountains. Nature 505, 43–49 (2014).

4. M. Meyer et al., A high-coverage genome sequence from an archaic Denisovan individual. Science 338, 222–226 (2012).

5. S. R. Browning, B. L. Browning, Y. Zhou, S. Tucci, J. M. Akey, Analysis of Human Sequence Data Reveals Two Pulses of Archaic Denisovan Admixture. Cell 173, 53–61 e59 (2018).

6. F. A. Villanea, J. G. Schraiber, Multiple episodes of interbreeding between Neanderthal and modern humans. Nat Ecol Evol 3, 39–44 (2019).

7. G. S. Jacobs et al., Multiple Deeply Divergent Denisovan Ancestries in Papuans. Cell 177, 1010–1021 e1032 (2019).

8. I. Juric, S. Aeschbacher, G. Coop, The Strength of Selection against Neanderthal Introgression. PLoS Genet 12, e1006340 (2016).

9. M. Petr, S. Paabo, J. Kelso, B. Vernot, Limits of long-term selection against Neandertal introgression. Proc Natl Acad Sci U S A 116, 1639–1644 (2019).

10. R. M. Gittelman et al., Archaic Hominin Admixture Facilitated Adaptation to Out-of-Africa Environments. Curr Biol 26, 3375–3382 (2016).

11. D. M. Vespasiani et al., Denisovan introgression has shaped the immune system of present-day Papuans. PLoS Genet 18, e1010470 (2022).

12. M. A. Almarri et al., Population Structure, Stratification, and Introgression of Human Structural Variation. Cell 182, 189–199 e115 (2020).

13. D. Enard, D. A. Petrov, Evidence that RNA Viruses Drove Adaptive Introgression between Neanderthals and Modern Humans. Cell 175, 360–371 e313 (2018).

14. P. Hsieh et al., Adaptive archaic introgression of copy number variants and the discovery of previously unknown human genes. Science 366, (2019).

15. E. Huerta-Sanchez et al., Altitude adaptation in Tibetans caused by introgression of Denisovan-like DNA. Nature 512, 194–197 (2014).

16. S. M. Yan et al., Local adaptation and archaic introgression shape global diversity at human structural variant loci. Elife 10, (2021).

17. P. Ebert et al., Haplotype-resolved diverse human genomes and integrated analysis of structural variation. Science 372, (2021).

18. G. A. Logsdon et al., The variation and evolution of complete human centromeres. Nature 629, 136–145 (2024).

19. H. Jeong et al., Structural polymorphism and diversity of human segmental duplications. Nat Genet 57, 390–401 (2025).

20. B. E. Morrow, D. M. McDonald-McGinn, B. S. Emanuel, J. R. Vermeesch, P. J. Scambler, Molecular genetics of 22q11.2 deletion syndrome. Am J Med Genet A 176, 2070–2081 (2018).

21. W. W. Liao et al., A draft human pangenome reference. Nature 617, 312–324 (2023).

22. P. Hsieh et al., Evidence for opposing selective forces operating on human-specific duplicated TCAF genes in Neanderthals and humans. Nat Commun 12, 5118 (2021).

23. M. Y. Dennis, E. E. Eichler, Human adaptation and evolution by segmental duplication. Curr Opin Genet Dev 41, 44–52 (2016).

24. M. R. Vollger et al., Segmental duplications and their variation in a complete human genome. Science 376, eabj6965 (2022).

25. S. Nurk et al., The complete sequence of a human genome. Science 376, 44–53 (2022).

26. B. Vernot et al., Excavating Neandertal and Denisovan DNA from the genomes of Melanesian individuals. Science 352, 235–239 (2016).

27. N. Brucato et al., Papua New Guinean Genomes Reveal the Complex Settlement of North Sahul. Mol Biol Evol 38, 5107–5121 (2021).

28. J. Adrian, P. Bonsignore, S. Hammer, T. Frickey, C. R. Hauck, Adaptation to Host-Specific Bacterial Pathogens Drives Rapid Evolution of a Human Innate Immune Receptor. Curr Biol 29, 616–630 e615 (2019).

29. M. Byrska-Bishop et al., High-coverage whole-genome sequencing of the expanded 1000 Genomes Project cohort including 602 trios. Cell 185, 3426–3440 e3419 (2022).

30. M. Rautiainen et al., Telomere-to-telomere assembly of diploid chromosomes with Verkko. Nat Biotechnol 41, 1474–1482 (2023).

31. D. Heller, M. Vingron, SVIM-asm: structural variant detection from haploid and diploid genome assemblies. Bioinformatics 36, 5519–5521 (2021).

32. M. Smolka et al., Detection of mosaic and population-level structural variants with Sniffles2. Nat Biotechnol 42, 1571–1580 (2024).

33. A. Paparella et al., Structural Variation Evolution at the 15q11-q13 Disease-Associated Locus. Int J Mol Sci 24, (2023).

34. Y. H. Zhou et al., Genetic Modifiers of Cystic Fibrosis Lung Disease Severity: Whole-Genome Analysis of 7,840 Patients. Am J Respir Crit Care Med 207, 1324–1333 (2023).

35. R. Hoftberger et al., Tubulin polymerization promoting protein (TPPP/p25) as a marker for oligodendroglial changes in multiple sclerosis. Glia 58, 1847–1857 (2010).

36. X. Chen et al., Association of nsv823469 copy number loss with decreased risk of chronic obstructive pulmonary disease and pulmonary function in Chinese. Sci Rep 7, 40060 (2017).

37. F. Yilmaz et al., Paleolithic Gene Duplications Primed Adaptive Evolution of Human Amylase Locus Upon Agriculture. bioRxiv, (2024).

38. D. Porubsky et al., Recurrent inversion polymorphisms in humans associate with genetic instability and genomic disorders. Cell 185, 1986–2005 e1926 (2022).

39. A. J. Sharp et al., A recurrent 15q13.3 microdeletion syndrome associated with mental retardation and seizures. Nat Genet 40, 322–328 (2008).

40. N. Kalebic et al., Human-specific ARHGAP11B induces hallmarks of neocortical expansion in developing ferret neocortex. Elife 7, (2018).

41. D. Yoo et al., Complete sequencing of ape genomes. Nature 641, 401–418 (2025).

42. F. Racimo et al., Archaic Adaptive Introgression in TBX15/WARS2. Mol Biol Evol 34, 509–524 (2017).

43. D. Yermakovich et al., Denisovan admixture facilitated environmental adaptation in Papua New Guinean populations. Proc Natl Acad Sci U S A 121, e2405889121 (2024).

44. G. A. Purnomo et al., Mitogenomes Reveal Two Major Influxes of Papuan Ancestry across Wallacea Following the Last Glacial Maximum and Austronesian Contact. Genes (Basel) 12, (2021).

45. S. Aganezov et al., A complete reference genome improves analysis of human genetic variation. Science 376, eabl3533 (2022).

46. F. A. Villanea et al., The MUC19 gene in Denisovans, Neanderthals, and Modern Humans: An Evolutionary History of Recurrent Introgression and Natural Selection. bioRxiv, (2024).

47. S. B. Cassidy, S. Schwartz, J. L. Miller, D. J. Driscoll, Prader-Willi syndrome. Genet Med 14, 10–26 (2012).

48. N. Brucato et al., Chronology of natural selection in Oceanian genomes. iScience 25, 104583 (2022).

49. J. Ebler et al., Pangenome-based genome inference allows efficient and accurate genotyping across a wide spectrum of variant classes. Nat Genet 54, 518–525 (2022).

50. D. Bueno, M. K. E. Schafer, S. Wang, M. J. Schmeisser, A. Methner, NECAB family of neuronal calcium-binding proteins in health and disease. Neural Regen Res 20, 1236–1243 (2025).

51. T. Ruan et al., Deficiency of IQCH causes male infertility in humans and mice. Elife 12, (2024).

52. L. X. Wang, M. R. Frey, R. Kohli, The Role of FGF19 and MALRD1 in Enterohepatic Bile Acid Signaling. Front Endocrinol (Lausanne) 12, 799648 (2021).

53. J. Yang et al., Integrative analysis of transcriptome-wide association study and gene expression profiling identifies candidate genes associated with stroke. PeerJ 7, e7435 (2019).

54. T. E. Thorgeirsson et al., Rare loss-of-function variants in HECTD2 and AKAP11 confer risk of bipolar disorder. Nat Genet 57, 851–855 (2025).

55. S. Yang, C. Zheng, C. Xia, J. Kang, L. Gu, Detection of positive selection on depression-associated genes. Heredity (Edinb) 134, 263–272 (2025).

56. K. Tretina, E. S. Park, A. Maminska, J. D. MacMicking, Interferon-induced guanylate-binding proteins: Guardians of host defense in health and disease. J Exp Med 216, 482–500 (2019).

57. E. P. Baker et al., Evolution of host-microbe cell adherence by receptor domain shuffling. Elife 11, (2022).

58. Y. Murata et al., Protein tyrosine phosphatase SAP-1 protects against colitis through regulation of CEACAM20 in the intestinal epithelium. Proc Natl Acad Sci U S A 112, E4264–4271 (2015).

59. Y. Souilmi et al., Admixture has obscured signals of historical hard sweeps in humans. Nat Ecol Evol 6, 2003–2015 (2022).

60. A. J. Stern, P. R. Wilton, R. Nielsen, An approximate full-likelihood method for inferring selection and allele frequency trajectories from DNA sequence data. PLoS Genet 15, e1008384 (2019).

61. B. Shaw et al., Emergence of a Neolithic in highland New Guinea by 5000 to 4000 years ago. Sci Adv 6, eaay4573 (2020).

62. S. H. Huang et al., Phage display technique identifies the interaction of severe acute respiratory syndrome coronavirus open reading frame 6 protein with nuclear pore complex interacting protein NPIPB3 in modulating Type I interferon antagonism. J Microbiol Immunol Infect 50, 277–285 (2017).

63. C. Bekpen, D. Tautz, Human core duplicon gene families: game changers or game players? Brief Funct Genomics 18, 402–411 (2019).

64. S. A. Langley, K. H. Miga, G. H. Karpen, C. H. Langley, Haplotypes spanning centromeric regions reveal persistence of large blocks of archaic DNA. Elife 8, (2019).

65. C. Posth et al., Deeply divergent archaic mitochondrial genome provides lower time boundary for African gene flow into Neanderthals. Nat Commun 8, 16046 (2017).

66. M. W. Nachman, S. L. Crowell, Estimate of the mutation rate per nucleotide in humans. Genetics 156, 297–304 (2000).

67. M. D. Kessler et al., De novo mutations across 1,465 diverse genomes reveal mutational insights and reductions in the Amish founder population. Proc Natl Acad Sci U S A 117, 2560–2569 (2020).

68. H. Jonsson et al., Parental influence on human germline de novo mutations in 1,548 trios from Iceland. Nature 549, 519–522 (2017).

69. D. Porubsky et al., Human de novo mutation rates from a four-generation pedigree reference. Nature, (2025).

70. T. P. Denham et al., Origins of agriculture at Kuk Swamp in the highlands of New Guinea. Science 301, 189–193 (2003).

71. L. Chmatal et al., Centromere strength provides the cell biological basis for meiotic drive and karyotype evolution in mice. Curr Biol 24, 2295–2300 (2014).

72. L. L. Sullivan, K. Chew, B. A. Sullivan, alpha satellite DNA variation and function of the human centromere. Nucleus 8, 331–339 (2017).

73. M. C. Zody et al., Evolutionary toggling of the MAPT 17q21.31 inversion region. Nat Genet 40, 1076–1083 (2008).

74. K. M. Steinberg et al., Structural diversity and African origin of the 17q21.31 inversion polymorphism. Nat Genet 44, 872–880 (2012).

75. M. R. Vollger et al., Long-read sequence and assembly of segmental duplications. Nat Methods 16, 88–94 (2019).

76. A. Rhie, B. P. Walenz, S. Koren, A. M. Phillippy, Merqury: reference-free quality, completeness, and phasing assessment for genome assemblies. Genome Biol 21, 245 (2020).

77. N. Huang, H. Li, compleasm: a faster and more accurate reimplementation of BUSCO. Bioinformatics 39, (2023).

78. J. A. Bailey, A. M. Yavor, H. F. Massa, B. J. Trask, E. E. Eichler, Segmental duplications: organization and impact within the current human genome project assembly. Genome Res 11, 1005–1017 (2001).

79. A. Seguin-Orlando et al., Paleogenomics. Genomic structure in Europeans dating back at least 36,200 years. Science 346, 1113–1118 (2014).

80. X. Yi et al., Sequencing of 50 human exomes reveals adaptation to high altitude. Science 329, 75–78 (2010).

81. M. Nei, W. H. Li, Mathematical model for studying genetic variation in terms of restriction endonucleases. Proc Natl Acad Sci U S A 76, 5269–5273 (1979).

82. S. H. Martin, J. W. Davey, C. D. Jiggins, Evaluating the use of ABBA-BABA statistics to locate introgressed loci. Mol Biol Evol 32, 244–257 (2015).

83. B. S. Weir, C. C. Cockerham, Estimating F-Statistics for the Analysis of Population Structure. Evolution 38, 1358–1370 (1984).

84. R. C. Lewontin, On measures of gametic disequilibrium. Genetics 120, 849–852 (1988).

85. F. Baumdicker et al., Efficient ancestry and mutation simulation with msprime 1.0. Genetics 220, (2022).

86. M. R. Vollger, P. Kerpedjiev, A. M. Phillippy, E. E. Eichler, StainedGlass: interactive visualization of massive tandem repeat structures with identity heatmaps. Bioinformatics 38, 2049–2051 (2022).

87. A. J. Drummond, G. K. Nicholls, A. G. Rodrigo, W. Solomon, Estimating mutation parameters, population history and genealogy simultaneously from temporally spaced sequence data. Genetics 161, 1307–1320 (2002).

88. M. Kimura, The neutral theory of molecular evolution. (Cambridge University Press, Cambridge Cambridgeshire ; New York, 1983), pp. xv, 367 p.

89. C. Jain, A. Rhie, N. F. Hansen, S. Koren, A. M. Phillippy, Long-read mapping to repetitive reference sequences using Winnowmap2. Nat Methods 19, 705–710 (2022).

90. H. Li, Minimap2: pairwise alignment for nucleotide sequences. Bioinformatics 34, 3094–3100 (2018).

91. A. Smit, Hubley, R & Green, P. (2015).

92. G. Benson, Tandem repeats finder: a program to analyze DNA sequences. Nucleic Acids Res 27, 573–580 (1999).

93. S. F. Altschul, W. Gish, W. Miller, E. W. Myers, D. J. Lipman, Basic local alignment search tool. J Mol Biol 215, 403–410 (1990).

94. R. S. Harris, The Pennsylvania State University (2007).

95. L. Skov et al., Detecting archaic introgression using an unadmixed outgroup. PLoS Genet 14, e1007641 (2018).

96. Y. Zhou, S. R. Browning, Protocol for detecting introgressed archaic variants with SPrime. STAR Protoc 2, 100550 (2021).

97. A. R. Quinlan, I. M. Hall, BEDTools: a flexible suite of utilities for comparing genomic features. Bioinformatics 26, 841–842 (2010).

98. K. Prufer, snpAD: an ancient DNA genotype caller. Bioinformatics 34, 4165–4171 (2018).

99. M. E. Lauterbur et al., Expanding the stdpopsim species catalog, and lessons learned for realistic genome simulations. Elife 12, (2023).

100. H. Li, R. Durbin, Inference of human population history from individual whole-genome sequences. Nature 475, 493–496 (2011).

101. S. Schiffels, K. Wang, MSMC and MSMC2: The Multiple Sequentially Markovian Coalescent. Methods Mol Biol 2090, 147–166 (2020).

102. F. Kumara Mastrorosa et al., Identification and annotation of centromeric hypomethylated regions with Centromere Dip Region (CDR)-Finder. bioRxiv, (2024).

103. J. A. Bailey et al., Recent segmental duplications in the human genome. Science 297, 1003–1007 (2002).

